# Loss of the Ap_4_A hydrolase YqeK impairs stress adaptation and virulence gene expression in *Staphylococcus aureus*

**DOI:** 10.64898/2026.05.18.725918

**Authors:** J. Vidaud, J.F. Coker, J. Silva, G.S. Davidson, C.C. Anderson, G.M. Bassett, A.A. Harry, T. Dusenbury, T.M. Gardner, M.L. Melear, N.J. Moraga, J.D. Fender, C.M. McMahon, M.R. Grosser

## Abstract

The nucleotide diadenosine tetraphosphate (Ap_4_A) accumulates during stress across organisms and cell types and is widely hypothesized to be an alarmone or second messenger. While Gram-negative bacteria use ApaH-family hydrolases to degrade Ap_4_A and other dinucleoside tetraphosphates (Ap_4_Ns), Gram-positive bacteria, including *Staphylococcus aureus,* use YqeK. Inactivation of Ap_4_A hydrolases and corresponding Ap_4_A accumulation cause diverse phenotypic effects in both Gram-negative and Gram-positive bacteria, ranging from increased sensitivity to antimicrobials to reduced virulence. However, the physiological role of YqeK in *S. aureus* remains uncharacterized. Here, we constructed an isogenic *yqeK* mutant in *S. aureus* and showed that Δ*yqeK* was sensitive to nitrosative and organic acid stress. We used a luminescence-based assay to show that Δ*yqeK* had ∼1000-fold higher relative Ap_4_N levels than wild-type even during unstressed growth, and all phenotypes were restored by complementation. Transcriptomics revealed that Δ*yqeK* exhibited stress-specific dysregulation of translation, nucleotide metabolism, central metabolism, iron acquisition, and stress response genes. In contrast, Δ*yqeK* had few transcriptional differences relative to wild-type during unstressed growth despite the large Ap_4_N accumulation, suggesting that the effects of Ap_4_Ns are contingent on the cellular stress state. Unexpectedly, we also found that the entire *agr* quorum sensing operon and numerous additional virulence genes, including hemolytic toxins, had reduced expression in Δ*yqeK*, correlating with reduced hemolytic activity in the mutant even in the absence of stress. Our data reveal YqeK to be a critical metabolic determinant of *S. aureus* stress resistance and virulence and position this hydrolase as a promising candidate for anti-virulence drug development.

**Importance:** *S. aureus* is a leading cause of antibiotic-resistant bacterial infections worldwide and is resistant to many components of the host immune response. Here, we discovered that deletion of YqeK, an enzyme that degrades a stress-associated nucleotide signaling molecule called Ap_4_A, rendered *S. aureus* more susceptible to infection-relevant stress conditions but had little impact on normal growth. Ap_4_Ns accumulated in the *yqeK* mutant and caused major stress-specific changes in gene expression, including reduced expression of key virulence genes. This correlated with a reduction in the destruction of red blood cells, a measure of bacterial toxicity toward host cells. Our data suggest that YqeK represents a promising target for new drugs aimed at reducing the virulence of *S. aureus*.

## Introduction

Nucleotide signaling molecules are widespread and diverse among prokaryotic and eukaryotic cells. The stress-associated nucleotide diadenosine tetraphosphate (Ap_4_A) was first discovered in the 1960s and is found across all kingdoms of life, from bacteria to mammals (1–3). It is primarily produced by aminoacyl-tRNA synthetases when the aminoacyl-AMP intermediate transfers AMP to ATP rather than a tRNA substrate (4). Aminoacyl-tRNA synthetases have varying capacity to perform this reaction, and other Ap_4_Ns and Np_n_Ns (where N is A, C, G, or U and n is the number of phosphates) are also produced (5, 6).

Across cell types, Ap_4_A levels generally increase during stress (3). After Ap_4_A accumulation was first observed during heat, cadmium, and oxidative stress in *Salmonella typhimurium* and *Escherichia coli* in the 1980s, a role for Ap_4_A as a possible alarmone, or stress signaling molecule, was proposed (7, 8). It has since been debated whether Ap_4_Ns have a signaling role or rather are stress or damage indicators (2, 3, 9, 10). In support of the alarmone hypothesis, in *Bacillus subtilis*, Ap_4_A binds and regulates activity of inosine-5’-monophosphate dehydrogenase (IMPDH), an enzyme in the guanine biosynthesis pathway (11). *B. subtilis* AcuB, a lysine deacetylase inhibitory protein, also binds and is stabilized by Ap_4_A, linking Ap_4_A to the regulation of protein acetylation (12). A recent proteomics study identified 87 putative Ap_4_A or Ap_3_A binding proteins in human embryonic kidney cells and 20 possible interactor proteins in *E. coli* (13). In addition to modulating protein function, *E. coli* and *B. subtilis* incorporate Ap_4_Ns as 5’ mRNA caps that reduce transcript digestion by exonucleases (14, 15).

Intracellular concentrations of Ap_4_A are regulated by the activity of dedicated Ap_4_A hydrolases. While mammalian cells use NUDIX-family enzymes for asymmetrical degradation of Ap_4_A (16), most Gram-negative bacteria primarily regulate Ap_4_A levels via ApaH family hydrolases that symmetrically hydrolyze Ap_4_A into two ADP (17). In contrast, Firmicutes, including *S. aureus*, lack ApaH family hydrolases and instead utilize YqeK, an HD-domain enzyme with a di-iron catalytic cluster, for symmetrical hydrolysis of Ap_4_A (18). ApaH and YqeK can also hydrolyze the 5’ Ap_4_N caps on mRNAs, exposing transcripts to exonuclease digestion (14, 15). The enzymatic function of *S. aureus* YqeK has been well-characterized *in vitro* and via ectopic expression in *B. subtilis* (18), but not yet in live *S. aureus* cells. Purified *S. aureus* YqeK can hydrolyze both Ap_4_A and other Ap_4_N nucleotides, but has high specificity for Ap_4_A.

Deletion of Ap_4_A hydrolases across bacterial species results in accumulation of Ap_4_A to levels ranging from 5 to 100-fold greater than those of wild-type (WT) (19–21) and is associated with fitness defects during specific stressors. This has been most well-characterized in Gram-negative species, where excess Ap_4_A accumulation exacerbates oxidative stress (21, 22), heat stress (21), and aminoglycoside sensitivity (23), as well as impacting virulence (20, 24), biofilm formation (25, 26), sporulation (19), and motility (22, 27). In *Pseudomonas aeruginosa*, for example, deletion of *apaH* results in attenuated virulence in plant and animal models due to reduced expression of virulence factors including quorum sensing signals, proteases, and siderophores (20). Although less well-studied, similar results have recently been reported in Gram-positive bacteria. Deletion of the native *yqeK* from *B. subtilis* results in accumulation of Ap_4_A and other Ap_4_Ns and causes increased sensitivity to disulfide stress (14, 18). In *Streptococcus mutans*, *yqeK* deletion caused Ap_4_A accumulation and reduced production of water-insoluble extracellular polysaccharides required for biofilm formation (25). To date, however, the physiological role of YqeK has not been studied in *S. aureus*.

*S. aureus* is a leading cause of both hospital-acquired and community-acquired infections and can cause a range of infection presentations including skin and soft tissue infections (SSTIs), endocarditis, osteomyelitis, and bacteremia (28–30). Stress resistance is a critical feature of *S. aureus* virulence, as it must respond to multiple host-derived stressors during infection, including nitric oxide (NO·) produced by phagocytes and low pH, organic acid-rich environments such as abscesses and the skin surface (31–33). In a transposon-sequencing screen in 2018, *S. aureus yqeK* was identified in a list of genes important for fitness during NO· stress and survival in a mouse skin abscess, but no follow up was performed on this particular hit (34). Here, we used phenotypic assays and transcriptomics to show that an isogenic *yqeK* mutant accumulated Ap_4_Ns, was sensitive to nitrosative and acid stress, and exhibited widespread stress-specific gene dysregulation. Notably, the *yqeK* mutant had broadly reduced expression of quorum sensing and virulence genes, including those encoding toxins, proteases, and immune evasion proteins, which correlated with reduced hemolytic activity. Altogether, our results provide the first physiological characterization of YqeK in *S. aureus* and indicate that YqeK is a critical enzyme that links *S. aureus* stress resistance and virulence.

## Results

### Δ*yqeK* is sensitive to NO· stress

We first sought to validate whether YqeK contributes to NO· resistance in *S. aureus*, as suggested by a prior transposon-sequencing screen of a pooled transposon mutant library (34). We created an in-frame clean deletion mutant of *yqeK* in the *S. aureus* community-associated MRSA strain LAC (35) via allelic exchange. We also complemented the mutation with a functional copy of *yqeK* on the plasmid pRMC2, which exhibits leaky expression from its tetracycline-inducible promoter. Additionally, we created a *yqeK* overexpression strain by transforming WT with pRMC2-yqeK, providing an extra copy of the gene.

Next, we investigated growth phenotypes of these strains during NO· stress and unstressed growth in TSB. The *yqeK* mutant grew similarly to WT during exponential phase in unstressed conditions and showed a slight growth defect primarily during the transition to stationary phase (∼10 hours) (Figure 1a). In contrast, following the addition of the NO· donor DETA/NO to cultures during early exponential growth, the mutant showed an immediate growth defect that continued for the duration of the experiment (Figure 1a). To compare the overall impact of NO· stress on the *yqeK* mutant relative to WT, we plotted the OD_650_ ratio of each strain (stressed/non-stressed) over the course of the growth curve. NO· had a significantly disproportionate effect on growth of the *yqeK* mutant (Figure 1b and 1c). Complementation of the mutant fully restored growth during NO· stress to WT levels even without the addition of an inducer, and there was no difference in growth for the WT strain harboring an extra copy of *yqeK*.

**Figure 1.**
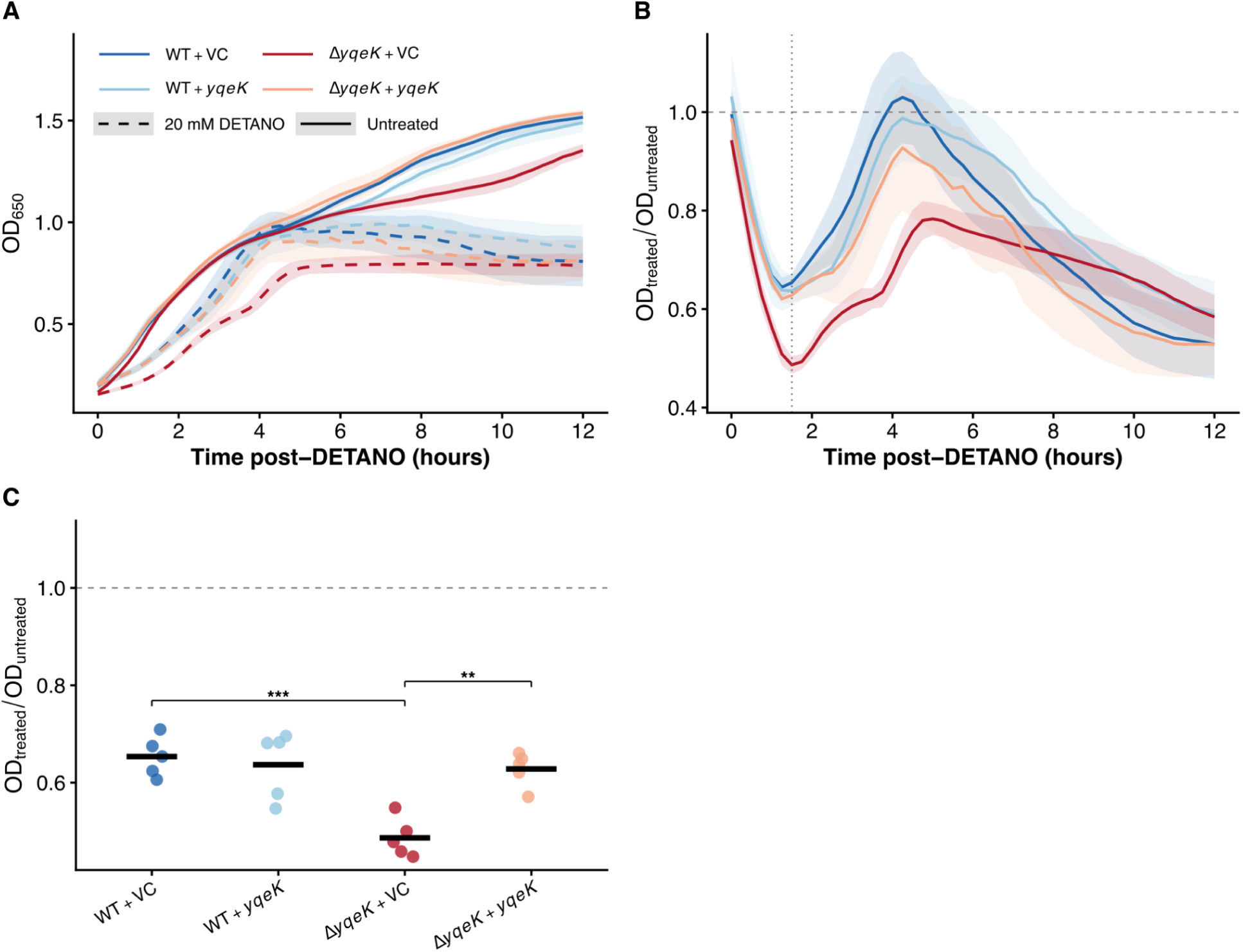
Δ*yqeK* is sensitive to NO• stress. Growth curves of the *S. aureus* LAC strains WT + pRMC2 (vector control, VC), Δ*yqeK* + VC, WT + pRMC2-*yqeK*, and Δ*yqeK* + pRMC2-*yqeK* (complemented mutant) were performed in 96-well plates in TSB with the addition of 20mM DETA/NO at the time point when OD_650_ was closest to 0.15. Plots depict growth of DETA/NO-exposed and control strains from the time of DETA/NO addition t=0. A) Growth curves are plotted as means; shaded regions represent SEM; solid lines= TSB only; dashed lines= DETA/NO-exposed. Growth curves are presented on a linear scale to facilitate direct comparison of OD₆₅₀ differences between strains, including the stationary phase growth defect observed in Δ*yqeK*. B) OD_650_ ratios of DETANO-treated to untreated wells are plotted as means; shaded regions represent SEM. A vertical dotted line indicates the nadir time point (1.5 h post DETA/NO addition) where the OD_650_ ratio for Δ*yqeK* + VC was at its minimum (i.e., peak NO· sensitivity). C) Dot plot of OD_650_ ratios at the nadir time point 1.5 h post-DETA/NO addition; crossbars indicate means. A one-way ANOVA followed by Tukey’s post hoc test was used to determine pairwise significance (*p < 0.05, **p < 0.01, ***p < 0.001, n=5 biological replicates).

### Δ*yqeK* exhibits a growth defect during organic acid stress

In other organisms, Ap_4_A accumulates in response to multiple stressors, and excessive concentrations can increase stress sensitivity; therefore, we next wanted to ask whether the Δ*yqeK* mutant exhibits stress sensitivity beyond NO·. We chose organic acid stress as an additional physiologically relevant stressor because it is common in abscesses and during skin colonization, and *S. aureus* acidifies its media via organic acid production during late growth in batch cultures (36), a time when we observed a slight growth defect for Δ*yqeK*. We found that the *yqeK* mutant exhibited a significantly reduced maximum growth rate relative to the WT and complemented strains during growth in TSB acidified to pH 5.5 with acetic acid (Figure 2). This phenotype was dose-dependent (Figure S1), and again the WT strain with an extra copy of *yqeK* did not differ phenotypically from WT.

**Figure 2.**
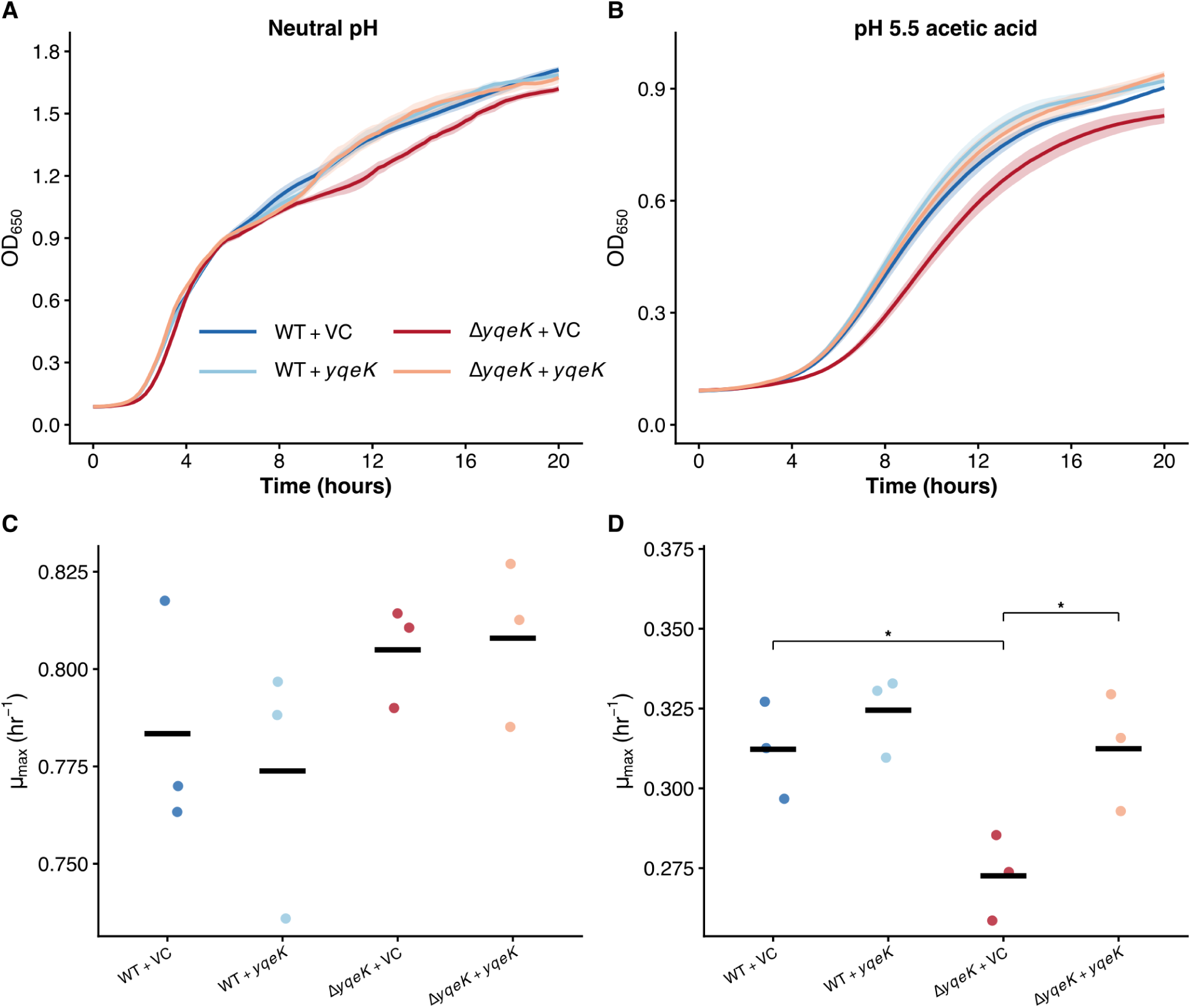
Δ*yqeK* exhibits a reduced growth rate during acetic acid stress. Growth curves of the *S. aureus* LAC strains WT + pRMC2 (vector control, VC), Δ*yqeK* + VC, WT + pRMC2-*yqeK*, and Δ*yqeK* + pRMC2-*yqeK* (complemented mutant) were performed in 96-well plates either in A) unmodified TSB (pH ∼7.3) or B) TSB acidified with acetic acid (pH 5.5). Growth curves are plotted as means; shaded regions represent SEM. Growth curves are presented on a linear scale to facilitate direct comparison of OD₆₅₀ differences between strains, including the stationary phase growth defect observed in Δ*yqeK*. Maximum specific growth rates (µ_max_) were calculated and plotted as dot plots for each strain during growth in C) TSB (pH ∼7.3) or D) acetic acid acidified TSB (pH 5.5) using the equation Δln OD_650_/Δtime for a sliding 2-h window. Crossbars indicate means. A one-way ANOVA followed by Tukey’s post hoc test was used to determine pairwise significance (*p < 0.05, n=3 biological replicates).

### ΔyqeK accumulates an excess of Ap_4_Ns

Although the enzymatic activity of purified *S. aureus* YqeK has been previously characterized (18), the cellular consequences of *yqeK* loss on Ap_4_A levels in native *S. aureus* has not yet been reported. To determine whether Ap_4_Ns accumulate in the *yqeK* mutant, we developed a luciferase-based assay to indirectly quantify Ap_4_Ns in each strain, based on the principles of an assay originally described by Ogilvie (37). We first depleted culture lysates of ATP via overnight incubation with a phosphatase. Then, a snake venom phosphodiesterase was used to hydrolyze Ap_4_Ns in the samples to NMP and ATP, which was quantified via luciferase activity on a plate reader. Relative luminescence units were more than 1000-fold greater in the *yqeK* mutant compared to those found in WT and complemented strain lysates (Figure 3). Moreover, during acid stress, there was a significant increase in RLUs in Δ*yqeK* relative to growth in TSB alone (Figure 3a), suggesting that Ap_4_N levels become further elevated during stress. RLUs for the WT and complemented strain were at the limit of detection for the assay, so we could not detect whether stress resulted in a slight increase in Ap_4_N concentrations in these strains. Altogether, these data confirm the physiological role of *S. aureus* YqeK as an Ap_4_N hydrolase.

**Figure 3.**
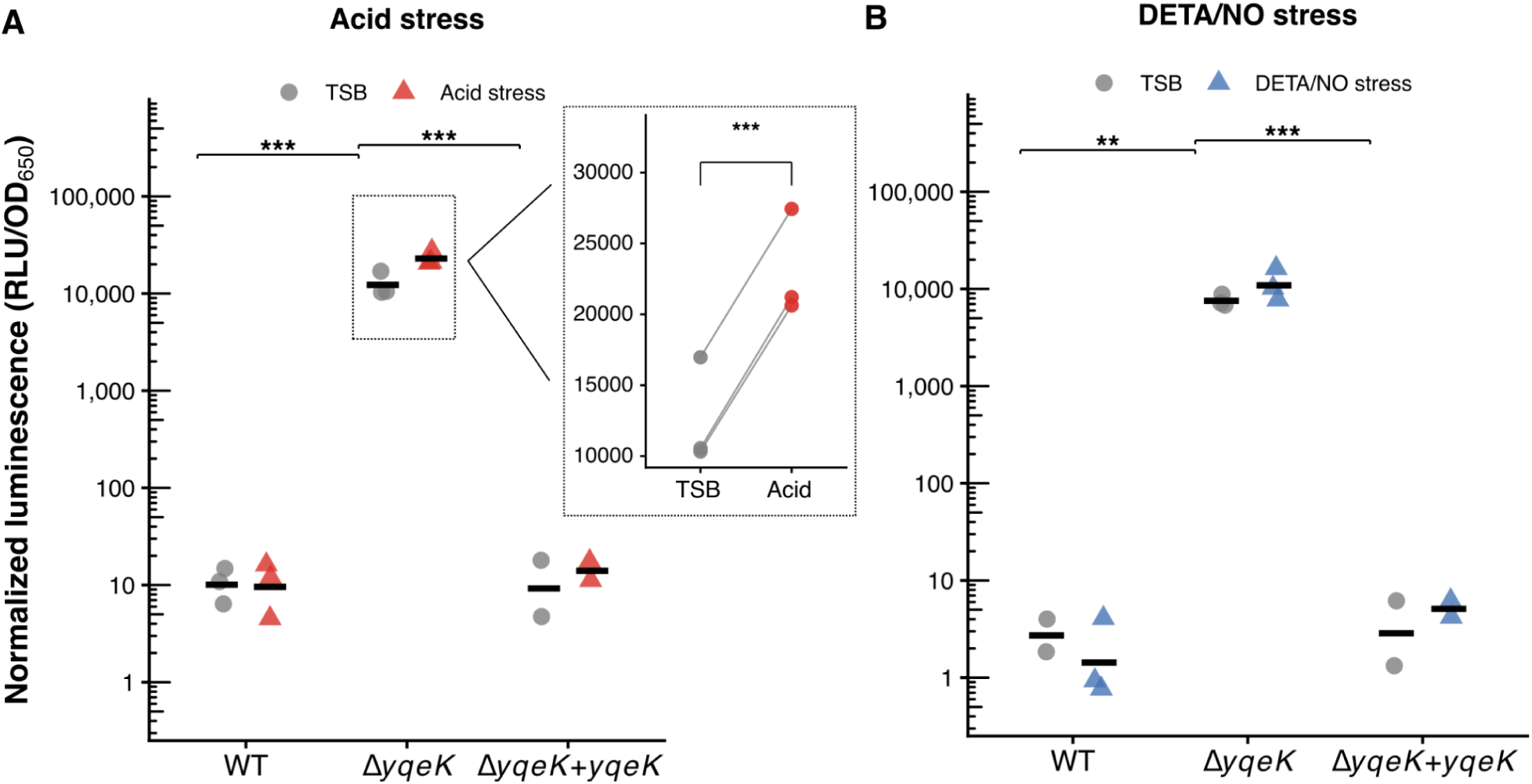
Ap_4_Ns accumulate in the Δ*yqeK* mutant under all growth conditions. Following ATP depletion of cell lysates for the strains WT, Δ*yqeK*, and Δ*yqeK* + pRMC2-*yqeK* (complemented strain), Ap_4_Ns were converted to ATP and AMP via treatment with snake venom phosphodiesterase. The resulting ATP was quantified via luciferase assay (BacTiter Glo) as an indirect measurement of relative Ap_4_Ns. A) Relative luminescence of Ap_4_Ns from overnight culture lysates during growth in TSB or acetic acid acidified TSB (pH 5.5) normalized to OD_650_. Inset shows pairwise comparisons for the Δ*yqeK* mutant in TSB control vs. acid stress. B) Relative luminescence units (RLUs), normalized to OD_650_, of Ap_4_Ns from lysates of log phase cultures following 1-hour exposure to 10mM DETA/NO (added at OD_650_ of 0.4), with control untreated strains (TSB only) grown in parallel. Crossbars indicate means. Significance was evaluated by two-way ANOVA on log-transformed normalized RLUs followed by Tukey’s post hoc test (asterisks indicate pairwise comparisons for the TSB control condition only; *p < 0.05, **p < 0.01, ***p < 0.001, n=3 biological replicates). In the inset, a planned paired t-test indicates that acid stress significantly elevated Ap_4_N levels in Δ*yqeK* relative to TSB control (p < 0.001).

### ΔyqeK exhibits widespread transcriptional dysregulation during stress

To investigate possible mechanisms for the Δ*yqeK* fitness defects during NO· and acid stress, we performed RNA-Seq on WT and Δ*yqeK* RNA extracted from cultures during unstressed growth in TSB, growth in TSB acidified to pH 5.5 with acetic acid, and growth following a 1-hour exposure to 10mM DETA/NO. We first examined the WT responses to acid stress and NO· stress relative to unstressed growth to validate our growth conditions. As expected, during acid stress, WT upregulated the urease operon and other genes typically associated with a response to acid stress (Figure S2, Table S2) (38). During NO· stress, WT induced expression of multiple genes typically involved in responding to nitrosative stress, including genes encoding anaerobic ribonucleotide reductases (*nrdD/nrdG*) and the flavohemoglobin Hmp (Figure S2, Table S2) (31).

We next compared transcription in Δ*yqeK* to WT within each condition to look for patterns that might explain the mutant’s growth defects. In TSB alone, only 28 genes were differentially expressed (Figure 4a). Conversely, during both stressors, Δ*yqeK* exhibited greater numbers of significantly altered genes (469 during acid stress, and 157 during NO· stress) (Figure 4b,4c). Translation machinery was one of the most prominent categories of genes upregulated in Δ*yqeK* under all three conditions (Figure 4). Top genes in this category included *tig*, encoding trigger factor, a ribosome-associated chaperone protein previously shown to be important during acid stress (39), and *gatCAB*, encoding a tRNA-dependent amidotransferase critical for proper tRNA charging (40).

**Figure 4.**
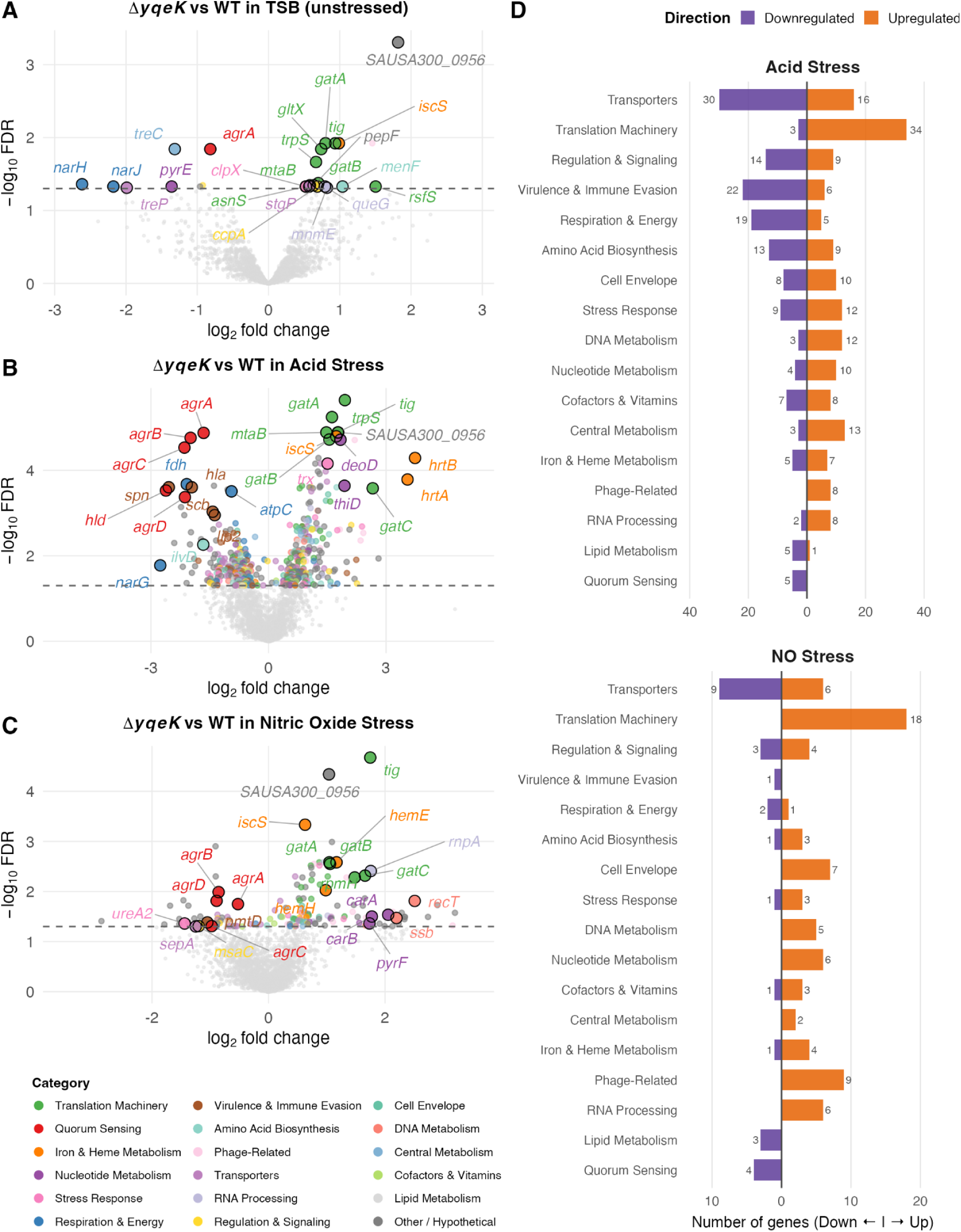
*S. aureus* Δ*yqeK* exhibits widespread gene dysregulation during acetic acid and NO· stress. Volcano plots of gene expression for the pairwise comparison of differential expression between Δ*yqeK* and WT during A) unstressed growth in TSB, B) acid stress and C) NO· stress. The TSB panel (A) is derived from the acid stress experimental batch. Genes were labeled based on a combined significance score (|log₂FC| × −log₁₀FDR) to highlight the most statistically robust differentially expressed genes. Labeled genes were selected from those with annotated functions; hypothetical proteins are plotted but not labeled for clarity. Select additional genes of biological relevance (e.g., pyrimidine biosynthesis, quorum sensing) that met the significance threshold (FDR < 0.05) are also labeled. Total significant genes: TSB, n = 28 (20 up, 8 down); acid stress, n = 469 (254 up, 215 down); NO· stress, n = 157 (112 up, 45 down). The *yqeK* gene was excluded from plots. D) The bar plot categorizes significant genes for each comparison during acid and NO· stress. Hypothetical or unannotated genes were excluded from the bar plot for clarity (246 and 68 genes for acid and NO· stress, respectively).

The gene SAUSA300_0956, encoding a predicted GNAT family N-acetyltransferase, was among the highest and most consistently upregulated genes in Δ*yqeK* across all three conditions (-log_10_FDR = 3.3, 4.9, and 4.3 in TSB, acid, and NO· stress respectively). Notably, a recent genome-wide association study reported that mutations in this gene were significantly associated with poor outcomes in *S. aureus* bacteremia, suggesting that it warrants further study for its potential role in pathogenesis (41).

Nucleotide metabolism genes were also aberrantly expressed in Δ*yqeK* with distinct patterns for each type of stress. During acid stress, Δ*yqeK* upregulated genes associated with purine interconversion (e.g., *deoD, xpt, guaB*) (Figure 4b). In contrast, during NO· stress, Δ*yqeK* failed to repress several pyrimidine biosynthesis genes (*carAB, pyrB, pyrC*) relative to WT (Figure 4c).

Other prominent categories of genes with dysregulated expression in the mutant during stress included central metabolism, iron acquisition, and stress-associated genes (Figure 4d, Figure 5). Genes associated with nitrate/nitrite respiration (e.g., *narH*) showed reduced expression in Δ*yqeK* under all conditions, including the unstressed TSB control. During acid stress in Δ*yqeK*, the entire ATP synthase operon showed reduced expression. Further, *hrtAB,* encoding an exporter of toxic free heme, was massively upregulated in Δ*yqeK,* and thioredoxin genes *trxA* and *trx* were also elevated. Heme biosynthesis genes (*hemE, hemH*) were upregulated under both stresses. WT significantly induced iron-acquisition genes (e.g., *sbn* operon, *isdA*, *sstA, feoB*) during NO· stress, but these genes were not significantly upregulated in Δ*yqeK*. Finally, lytic genes associated with the prophage φSa3 (encoding capsid, terminase, and connector proteins) were upregulated in Δ*yqeK* during acid stress.

**Figure 5.**
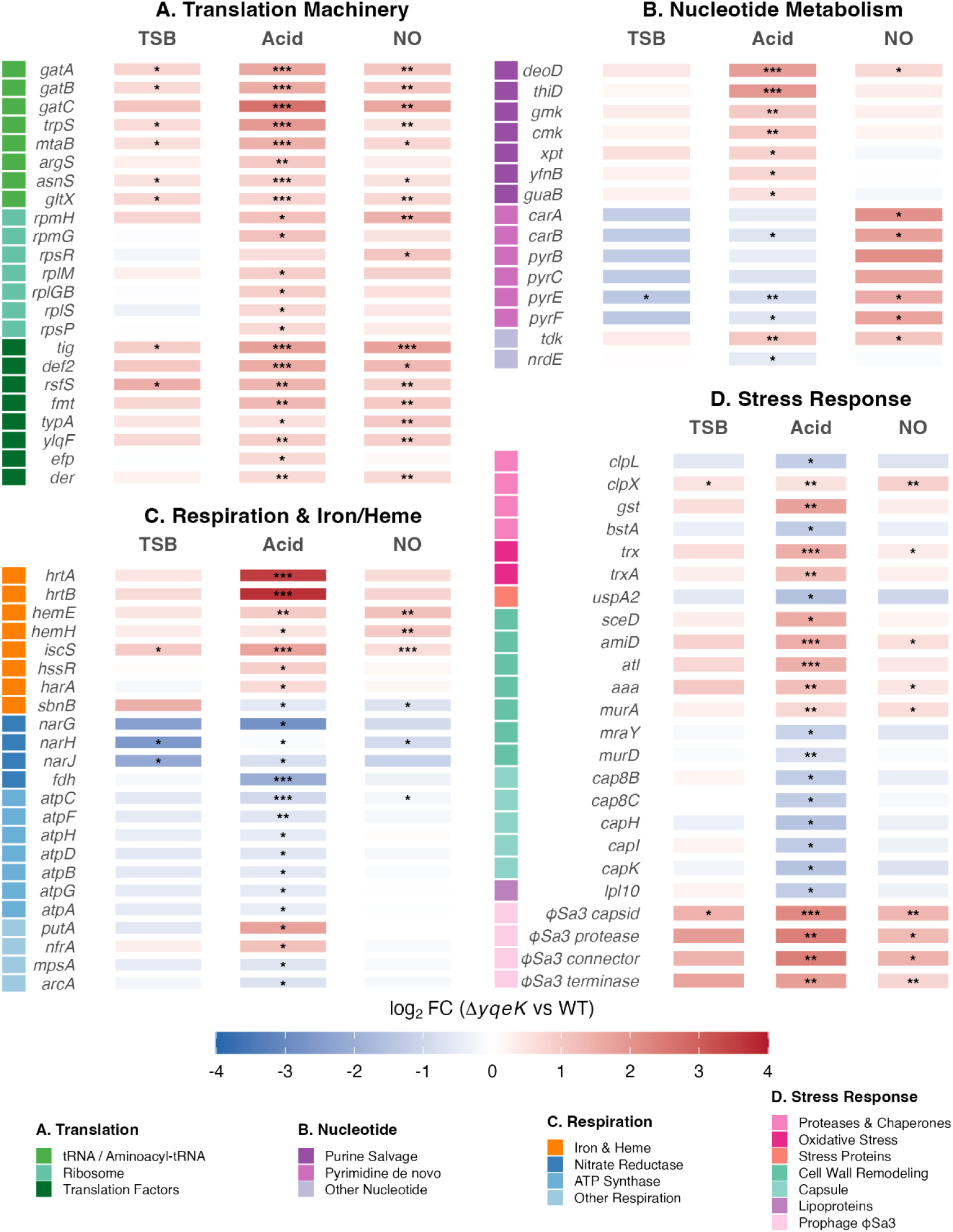
Major categories of genes with dysregulated gene expression in Δ*yqeK*. Heatmaps depict log_2_FC of genes associated with A) translation machinery, B) nucleotide metabolism, C) respiration and iron/heme, and D) stress response from RNA-Seq under unstressed (TSB), acetic acid stress, and NO· stress conditions. Genes are grouped and color-coded by function (sidebars). Blue indicates downregulation and red indicates upregulation in the Δ*yqeK* mutant relative to WT. Statistical significance is indicated by asterisks (*, FDR <0.05; **, FDR <0.01; ***, FDR <0.001).

Unexpectedly, RNA-seq results revealed an extensive reduction in virulence gene expression in Δ*yqeK* (Figure 7a). During both stressors, the full *agr* quorum sensing operon was significantly downregulated. Additional virulence factor genes showing reduced expression during acid stress included virulence regulators (*sarA, sae*), toxins (*hld* and *hla)*, immune evasion (e.g., *scn, scb*), secretion systems (*essB, essC*), surface and adhesion proteins, and enzymes (proteases, lipases). A similar trend was observed across these genes in the NO·-stressed samples, though fewer genes reached significance (Figure 7a). The gene encoding protein A (*spa)* is negatively regulated by *agr* (42), and, consistent with the *agr* down-regulation, we observed *spa* to be slightly upregulated in the Δ*yqeK* mutant.

**Figure 6.**
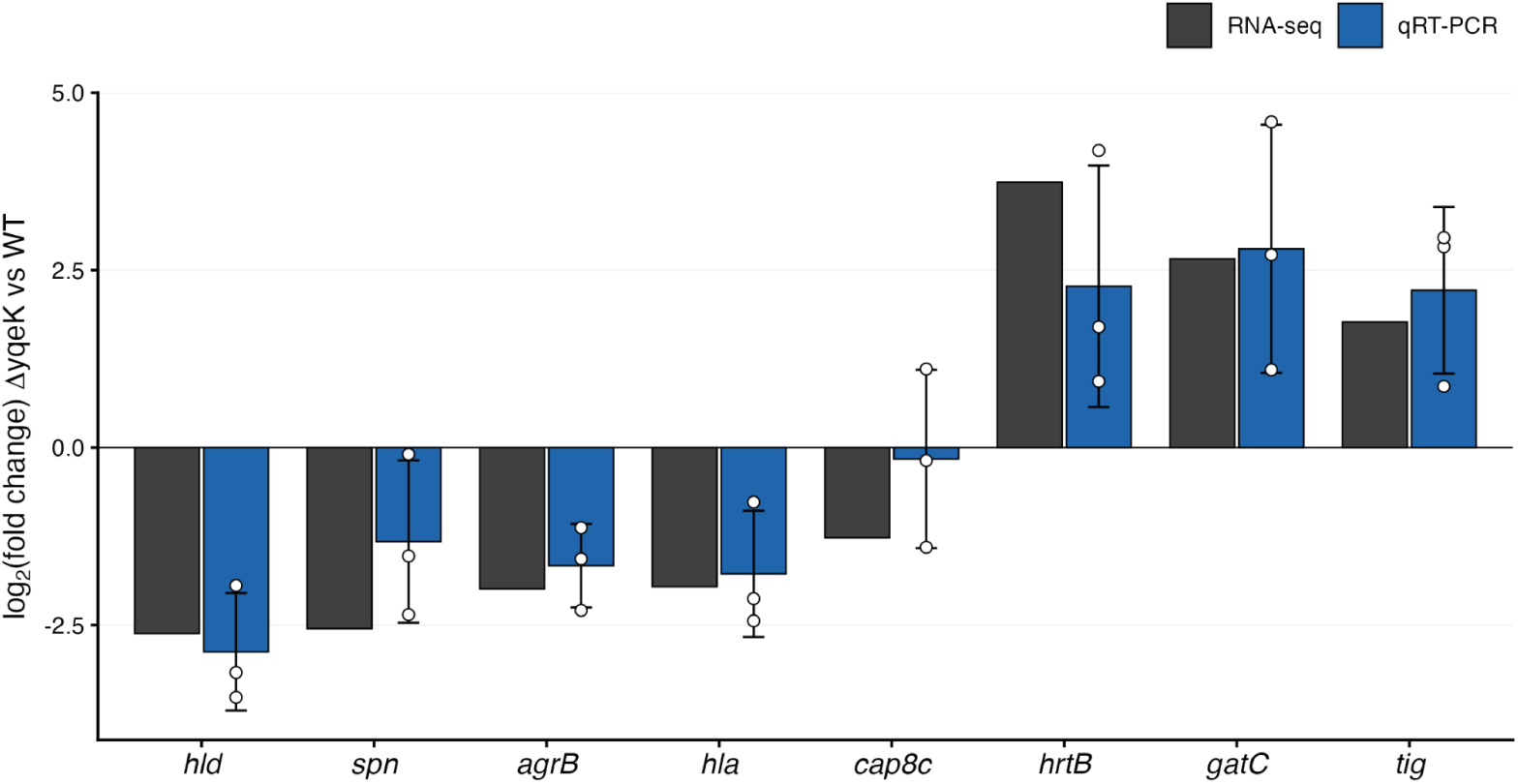
qRT-PCR validation of RNA-Seq results for acid stress experiment. qRT-PCR was performed on RNA collected from three additional biological replicates of WT and Δ*yqeK* in unstressed TSB and acetic acid stress conditions for genes in major categories: virulence (*hld, spn, agrB, hla*), stress response/capsule biosynthesis (*cap8c*), iron/heme metabolism (*hrtB*), and translation (*gatC, tig*). Log_2_ fold changes were calculated using the ΔΔCt method with normalization to the housekeeping gene *rpoD*. Bars show means +/- standard deviation; white dots show individual biological replicates. There was directional concordance for all three replicates for 7/8 genes, and for two of three replicates for *cap8c*, which had the smallest effect size. qRT-PCR fold changes correlated strongly with RNA-Seq fold changes (Pearson r = 0.950, p = 3.0 × 10⁻⁴; n=3 biological replicates).

**Figure 7.**
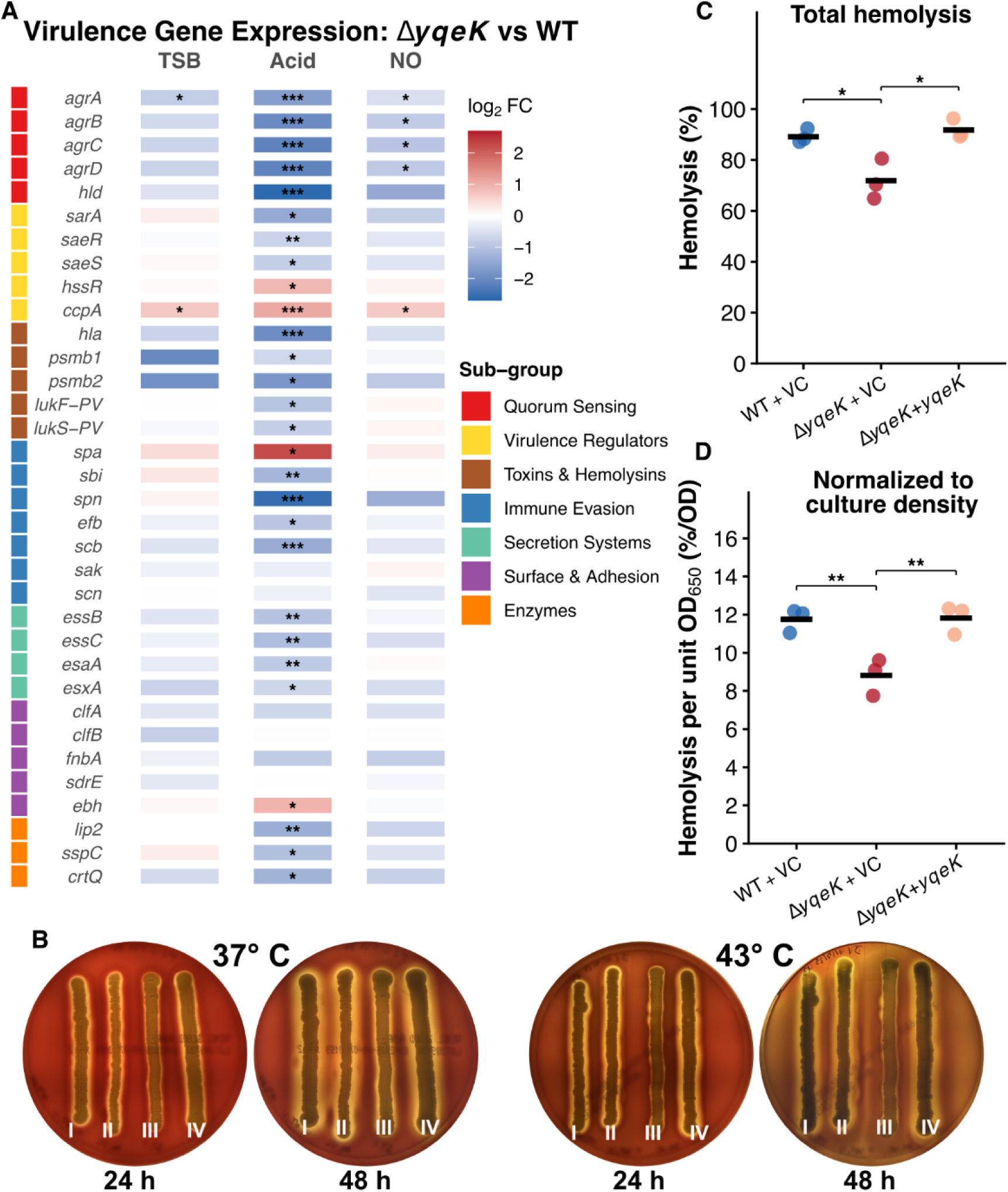
Δ*yqeK* has reduced virulence gene expression and hemolytic activity. A) Heatmap depicts log_2_FC of virulence-associated genes from RNA-Seq under unstressed (TSB), acetic acid stress, and NO· stress conditions. Genes are grouped and color-coded by function (sidebar). Blue indicates downregulation and red indicates upregulation in the Δ*yqeK* mutant relative to WT. Statistical significance is indicated by asterisks (*, FDR <0.05; **, FDR <0.01; ***, FDR <0.001). B) A qualitative hemolysis assay was performed on 5% sheep blood TSA plates grown at 37°C or 43°C and photographed at 24 and 48 hours; I) WT + VC, II) WT + *yqeK*, III) Δ*yqeK* + VC, IV) Δ*yqeK* + *yqeK*. C) and D) A quantitative liquid hemolysis assay was performed using 2% sheep erythrocytes incubated with filtered *S. aureus* overnight culture (TSB, 37°C, unstressed) supernatants for 1-hour. Remaining RBCs were pelleted, and hemoglobin release into supernatants was quantified on a plate reader by reading absorbance at 540nm. The % hemolysis was calculated as (sample - negative control)/(positive control - negative control)*100. Dot plots show total hemolysis (C) and OD-normalized hemolysis (D); crossbars show means. A one-way ANOVA followed by Tukey’s post hoc test was used to determine pairwise significance (*p < 0.05, **p<0.01; n=3 biological replicates).

To validate the RNA-Seq results, we collected new RNA samples from three additional biological replicates of WT and Δ*yqeK* during growth in regular and acidified TSB. We performed qRT-PCR on eight genes selected from major categories of dysregulated genes: virulence (*hld, spn, agrB, hla*), iron/heme metabolism (*hrtB*), stress response/capsule biosynthesis (*cap8c*), and translation (*tig*, *gatC*). The qRT-PCR fold changes were strongly correlated with RNA-Seq fold changes (Pearson r=0.950, p=3.0×10^-4^), and 7 of 8 genes showed directional concordance in all three qRT-PCR replicates (Figure 6). The exception was *cap8c*, which showed directional concordance in two of three replicates. This partial concordance likely reflects the relatively lower fold change of *cap8c* in the RNA-Seq (log_2_FC=-1.27), which may be at the limit of detection with n=3 biological replicates.

### Δ*yqeK* exhibits a hemolytic defect

To determine whether the lower expression of *agr*-regulated toxin genes in Δ*yqeK* translated to a reduction in toxin production, we investigated the hemolytic capability of the mutant relative to WT. First, we performed qualitative hemolysis assays on 5% sheep’s blood tryptic soy agar at 37°C and 43°C; the elevated temperature condition was included because heat stress is associated with Ap_4_A accumulation in other species (7). Δy*qeK* produced a notably smaller zone of hemolysis relative to the WT and complemented strains, and this difference was more pronounced on the 43°C plates (Figure 7b). Next, we performed a quantitative liquid hemolysis assay with sheep red blood cells using supernatants from overnight cultures of each strain. Overnights were cultured to similar optical densities at 37°C without stress to avoid possibly confounding growth and stress response differences between WT and Δ*yqeK*. The *yqeK* mutant exhibited ∼20% reduced hemolysis relative to WT and complemented strains even in the absence of stress (Figure 7c). Normalization to OD_650_ further increased this difference to ∼25% (Figure 7d), suggesting that the reduction is not attributable to differences in culture density and is consistent with the reduced expression of hemolytic toxin genes observed by RNA-Seq.

## Discussion

This study provides the first physiological characterization of an *S. aureus yqeK* deletion mutant and indicates that YqeK is critical for stress resistance and virulence in *S. aureus*. The markedly high levels of Ap_4_Ns in the Δ*yqeK* mutant (∼1000-fold greater than WT under multiple conditions tested) indicate that YqeK likely functions as the sole enzyme responsible for Ap_4_N clearance in *S. aureus*, with no redundant hydrolases able to compensate for its loss. A limitation of our luminescence-based assay is that it cannot distinguish between Ap_4_A and other Ap_4_Ns. Based on the catalytic specificity of purified YqeK for Ap_4_A *in vitro*, it is likely that Ap_4_A would accumulate disproportionately, although other Ap_4_Ns are also expected to accumulate (6, 18). Future LC-MS/MS studies could clarify the relative abundances of nucleotide species that accumulate.

Despite the large cytoplasmic concentration of Ap_4_Ns in Δ*yqeK* under all conditions, the *yqeK* mutant did not exhibit a substantial growth defect during growth in TSB until the transition to stationary phase, where it lagged behind WT before approaching WT levels by the end of the growth curve. In contrast, the more pronounced growth defect of Δ*yqeK* during NO· and acid stress suggests that excess intracellular Ap_4_N accumulation is disproportionately detrimental during stress.

Interestingly, RNA-Seq revealed relatively few transcriptional changes (only 28 genes) in the Δ*yqeK* mutant relative to WT during unstressed growth. Conversely, during either acid or NO· stress, hundreds of genes (469 and 157 respectively) were dysregulated. This striking stress-contingent pattern is consistent with Ap_4_Ns acting as *bona fide* alarmones in *S. aureus*, with effects that are activated by the cellular stress state and not by nucleotide concentration alone.

Our finding that nucleotide metabolism genes are significantly dysregulated in Δ*yqeK* suggests that Ap_4_N accumulation could be disrupting nucleotide pools, possibly by sequestering adenylate equivalents. In a recent study of a *P. aeruginosa apaH* mutant, quantification of nucleotide pools under unstressed conditions showed ∼20% reduction in ADP levels and no change in ATP, GTP, or GDP concentrations (20); however, it is possible that stress conditions may exacerbate changes to nucleotide pools or that the situation differs in *S. aureus*. In *B. subtilis*, the inhibition of IMPDH by Ap_4_A establishes a direct link between Ap_4_A and nucleotide metabolism, and suggests a possible mechanism by which Ap_4_A accumulation could be disrupting guanine nucleotide biosynthesis (11). Consistent with stress-induced purine pool disruptions, our RNA-Seq data showed upregulation of purine salvage genes during acid stress, including *deoD* (purine nucleoside phosphorylase), which liberates purine bases, as well as genes that could repurpose the liberated purines, such as *xpt* (xanthine phosphoribosyltransferase) and *guaB* (IMPDH, the Ap_4_A regulated enzyme in *B. subtilis*). During NO· stress, WT repressed *de novo* pyrimidine biosynthesis, but Δ*yqeK* failed to do so and even upregulated several of these genes (e.g., *carAB*), suggesting disruption of normal nucleotide pool sensing. These divergent patterns under acid and NO· stress reinforce that *yqeK* deletion causes stress-specific responses rather than a uniform disruption of nucleotide metabolism. Future metabolomics work could shed light on these hypotheses.

Beyond nucleotide pool disruption, the overall upregulation of translation-associated genes suggests that the excess of Ap_4_Ns is impairing translation and that the cell is compensating by increasing expression of aminoacyl-tRNA synthetases (ARS), ribosome assembly, and translation quality control genes. Disruption of GTP pools via IMPDH inhibition could affect the activity of translation elongation factors such as EF-Tu (43). Ap_4_A accumulation could reduce ATP pools via sequestration of adenylate equivalents, or could result in feedback inhibition of ARS activity, a mechanism demonstrated *in vitro* at supraphysiological Ap_4_A levels (44). Either mechanism could reduce the pool of charged tRNAs and lead to stalled ribosomes or translation fidelity issues; whether Ap_4_A concentrations reach sufficient levels in Δ*yqeK* to achieve ARS inhibition remains to be determined. The compensatory upregulation of ARS genes could lead to a possible positive feedback loop during stress, resulting in additional Ap_4_A production and exacerbating the problem.

In addition to aberrant nucleotide metabolism and translation, disruption of central metabolism and iron-acquisition pathways may also contribute to the severity of stress-related growth defects. The reduced expression of ATP synthase and anaerobic respiration genes in Δ*yqeK* suggests that energy generation may be impaired at multiple levels. During NO· stress, the induction of siderophore biosynthesis, heme acquisition, and iron transporter genes in WT is consistent with a need to repair damaged iron-sulfur clusters and restore iron homeostasis (31, 45). However, Δ*yqeK* fails to significantly induce these genes, which may lead to downstream impairment of iron-dependent respiratory and metabolic enzymes. This could reduce energy generation capabilities in Δ*yqeK* during NO· stress and may contribute to the mutant’s reduced growth phenotype.

There are several indicators that Δ*yqeK* may be experiencing greater oxidative and genotoxic stress than WT under comparable conditions. The upregulation of *hrtAB* and thioredoxin genes in Δ*yqeK* during acid stress (and upward trend in NO· stress) suggests heightened oxidative stress relative to WT, consistent with heme release from damaged proteins during these stressors (46). Additionally, the expression of the φSa3 prophage lytic genes during acid stress in Δ*yqeK* suggests genotoxic stress, perhaps related to issues with nucleotide metabolism.

Our most striking and unexpected finding from the RNA-seq data was the downregulation of the *agr* operon, along with the broad downregulation of additional virulence genes, under OD-matched mid-log conditions. These expression patterns were phenotypically validated by the reduced hemolytic activity of Δ*yqeK* supernatants, independently of NO· or acid stress. The *agr* quorum sensing system is among the most well-studied virulence regulators in *S. aureus*; the finding that YqeK loss suppresses its activity indicates that Ap_4_N signaling represents a previously unappreciated layer of *agr* regulation. The mechanism for reduced *agr* activity may be direct, through Ap_4_N interaction with components of the signaling pathway, or indirect, mediated either by other regulators or disruption of cellular energy levels. For example, AgrC has relatively low ATP affinity for a histidine kinase and is therefore particularly sensitive to reductions in cellular energy levels (47, 48); because *agr* is auto-regulated, even modest reductions in AgrC activity could be self-amplifying. Additionally, *sarA*, encoding a positive regulator of the *agr* system (49), is downregulated in the Δ*yqeK* mutant, suggesting possible indirect regulatory contributions. Collectively, these data suggest that inactivation of *yqeK* reduces virulence factor production and may attenuate pathogenesis *in vivo*, a hypothesis that future animal model studies will be needed to test. Importantly, the connection between Ap_4_Ns and virulence regulation may apply beyond *S. aureus*, since recent studies have linked Ap_4_A to virulence in *P. aeruginosa* and *S. mutans* (20, 25). Future studies should also investigate SAUSA300_0956, the GNAT family N-acetyltransferase consistently upregulated in Δ*yqeK* across all conditions and recently associated with poor *S. aureus* bacteremia outcomes (41), as a potential Ap_4_N-regulated pathogenesis determinant.

Taken together, our data implicate YqeK as a non-redundant Ap_4_N hydrolase at the intersection of metabolism, stress response, and virulence regulation in *S. aureus*. Our transcriptomics data indicate both shared and divergent transcriptional responses of Δ*yqeK* to acid and nitrosative stress, suggesting that Ap_4_Ns may have stress-specific functions during host-relevant stress conditions. The finding that Ap_4_N accumulation only minimally impacts unstressed mid-log cells despite dramatic effects during stress will require further mechanistic exploration; it is possible that stress may activate specific Ap_4_A-responsive pathways or that already stressed cells are more sensitized to the consequences of Ap_4_A accumulation. Future studies should investigate the relative contributions of nucleotide pool disruption, Ap_4_A protein binding partners, and/or 5’ mRNA capping by Ap_4_A to the phenotypic and transcriptional consequences of *yqeK* deletion. Because the YqeK hydrolase is unique to Gram-positive bacteria and does not significantly impact unstressed growth in *S. aureus*, it represents a promising anti-virulence drug target that would impose limited pressure for resistance development during normal growth, while possibly attenuating virulence *in vivo*.

## Materials and Methods

### Strains

Strains, plasmids, and primers are listed in Table S1. *S. aureus* strains were cultured in tryptic soy broth (TSB). For antibiotic selection of plasmids, 10µg/ml chloramphenicol was used. *S. aureus* USA300 strain LAC (35) was used for all phenotypic and transcriptomics experiments. A clean, in-frame deletion of *yqeK* (SAUSA300_1552) was created using the *E. coli* shuttle vector pBT2ts (50) for allelic exchange according to previously described methods (51). To create the knockout plasmid, the 5’ and 3’ flanking regions of *yqeK*, including the start and stop codons, were PCR amplified using primers p314, p315, p316, and p317 (Table S1). The fragments were then joined into a single fragment using a SOE PCR and cloned into the BamHI site of pBT2ts. For complementation, *yqeK* was amplified from LAC genomic DNA with primers p360 and p318. Because *yqeK* is located mid-operon, a strong Shine-Dalgarno sequence (TACCCGGAGGAGATAT) was added. This fragment was cloned into the tetracycline-inducible expression vector pRMC2 (Addgene plasmid #68940, a gift from Tim Foster; (52). Full complementation of the Δ*yqeK* mutant was achieved under most conditions without induction, due to leaky expression from the promoter, so no inducer was used. For growth curves and hemolysis experiments, all strains carried the pRMC2 plasmid (vector control or pRMC2_yqeK) to ensure equivalent antibiotic and metabolic pressures. For RNA-Seq and luminescence assays, we used WT and Δ*yqeK* strains without plasmids to avoid potentially confounding transcriptional effects of plasmid maintenance.

### Growth Curves

Growth assays were performed in a 96-well plate format with 200µL culture volumes. For inoculation, bacteria from overnight TSB cultures were diluted to an OD_650_ of 0.01. Growth was assessed via OD_650_ measurements taken every 15 minutes for 24 hours using a BioTek Synergy HTX plate reader set at 37°C with linear shaking.

For acid stress experiments, TSB (pH ∼7.3) was acidified to a pH of 5, 5.5, or 6 with acetic acid, and the acidified media was used when preparing the growth curve inoculum. Maximum specific growth rates (µmax) were calculated for strains in unmodified and acidified TSB using the equation Δln OD_650_/Δtime (in hours) for a sliding two hour window. Growth curve OD_650_ values were plotted on a linear scale to allow better visualization of differences in OD_650_ at all growth phases.

For nitric oxide stress experiments, the NO· donor 2,2′-(hydroxynitrosohydrazono)bis-ethanimine (DETA)/NO (Fisher Scientific, Waltham, MA) diluted in 0.01N NaOH was added at a concentration of 20mM at the time point when cultures were nearest an OD650 0.15. To control for small differences in OD_650_ at the time of treatment, NO· sensitivity was quantified and plotted as the ratio of treated to untreated OD_650_ for each strain for the remainder of the growth curve. The minimum ratio of treated/untreated for each strain (nadir ratio), corresponding to the time point at which growth was most affected by NO· stress (1.5 hours post DETA/NO addition), was compared between strains for statistical analysis.

### Ap_4_N Luminescence Assay

To quantify Ap_4_Ns in culture lysates, we developed a snake venom phosphodiesterase luciferase assay based on the principles of an assay originally described by Ogilvie (37). Stationary phase Ap_4_N concentrations were quantified from strains grown in TSB or TSB pH 5.5 with acetic acid. Overnight cultures were normalized to an OD_650_ of 5. Acid-grown cultures did not reach an OD_650_ of 5 and had to be concentrated by centrifugation and resuspension. Next, 1mL of normalized culture was pelleted at 21,300xg for 1-min, washed once with 1 mL phosphate buffered saline (PBS, pH 8), and resuspended in 1 mL Tris buffer (10mM, pH 8). Separately, late exponential phase Ap_4_N concentrations were quantified from cultures exposed to NO·. For these cultures, overnights of each strain were back-diluted 1:100 into fresh TSB, grown to an OD_650_ of 0.4, and exposed to 10mM DETA/NO for 1-hour. Cultures not exposed to NO· were grown simultaneously for comparison. After 1-hour, 5mL of each culture was pelleted, washed, and resuspended in Tris buffer as described above. For these experiments, we used plasmid-free WT and Δ*yqeK* strains, alongside the complemented Δ*yqeK* strain to verify that accumulation is specifically due to the absence of the *yqeK* gene.

Cells were lysed via treatment with 62.5µg/mL lysostaphin for 15-min at 37°C, followed by bead-beating at setting 5 for three cycles of 30 seconds each, with 2 minute incubations on ice between cycles. Lysates were centrifuged at 17,700 x g for 5 minutes to pellet beads and cell debris. The clarified supernatant (600µL) was transferred to a clean microtube and boiled at 95°C for 10 minutes to inactivate enzymes. Samples were centrifuged again at 21,300 x g for 2 minutes, and the supernatant was transferred to a clean microtube.

To deplete ATP from the lysates, samples were incubated overnight at 37°C with ∼25 units of Antarctic phosphatase (New England Biolabs), with gentle shaking at 150 rpm. Following incubation, the phosphatase was inactivated at 80°C for 5 minutes. Half of each lysate was reserved as a control to assess residual ATP, and the other half (∼300µL) was treated with 160µg/mL snake venom phosphodiesterase (Phosphodiesterase I from *Crotalus atrox*, Sigma Aldrich, Burlington, MA) for 10-min at room temperature to convert Ap_4_Ns to ATP and AMP. The resulting ATP was then quantified using the BacTiter-Glo Microbial Cell Viability Assay (Promega) as an indirect measure of Ap₄N levels. Briefly, 50 µL of each sample was added to 50 µL BacTiter-Glo reagent in a white-walled 96 well plate in triplicate, with adjacent wells left empty to minimize signal crosstalk. Luminescence was measured immediately following reagent addition (BioTek Synergy HTX). Relative luminescence units (RLU) of PDE-treated samples were normalized by subtracting background readings of sample lysates untreated with PDE (to account for any residual ATP) and then divided by OD_650_ to calculate RLU/OD_650_.

### RNA Collection

For RNA collection experiments, overnight cultures of each strain were back-diluted 1:100 into fresh media and grown to mid-exponential phase as follows. For the TSB control group and the acid treatment group, exponential cultures were grown in unmodified TSB or acidified TSB (pH 5.5 with acetic acid) to an OD_650_ of 0.5 and then were harvested. For the nitric oxide treatment group, cultures were grown in unmodified TSB as described above to an OD_650_ of 0.4, at which point 10 mM DETA/NO was added (a lower concentration than that used in growth curve experiments, due to the large culture volume required for RNA collection). Cultures were incubated for 1 hour at 37°C with shaking, then harvested. Cells were pelleted by centrifugation at 4°C for 5 minutes at 10,000 × g. The supernatant was removed and pellets were flash frozen in liquid nitrogen, then stored at-80°C until extraction. RNA was extracted using a Qiagen RNeasy kit following the modified protocol described in (53). Following extraction, DNA was removed via treatment with Turbo DNase (Invitrogen) for one hour. The TSB control RNA was prepared once from each biological replicate and sent twice for library prep and sequencing (once each with the acid and NO· experimental groups), so that independent libraries were generated from the same RNA in each experimental batch.

### RNA-Sequencing

Library prep and sequencing was performed by SeqCenter, LLC (Pittsburgh, PA). An additional DNase treatment was performed with Invitrogen DNase. Libraries were prepared using Illumina’s Stranded Total RNA Prep Ligation with Ribo-Zero Plus kit and 10bp unique dual indices (UDI). Libraries were sequenced on a NovaSeq X Plus to generate paired end 150bp reads. Demultiplexing, quality control, and adapter trimming was performed with bcl-convert (v4.2.4).

Read quality was assessed using FastQC (v0.12.1) through KBase. No further trimming was performed. Reads were aligned to reference genome *Staphylococcus aureus* subsp. aureus USA300_FPR3757 (NCBI RefSeq <u>NC_007793.1</u>) using HISAT2 (v2.1.0) with default parameters via the KBase platform. Read counting was performed in R using Subread featureCounts (v2.24.0) to generate a raw count matrix. Acid stress and NO· stress were separately analyzed as distinct experiments, each with its own independently sequenced TSB control libraries. Differential expression analysis was performed with edgeR (v4.8.2) using a factorial model ∼genotype * condition (genotype= WT or ΔyqeK; condition= control or stress; interaction= genotype:condition). TMM was used for normalization, and the data were fitted to a GLM model with factorial design, after which a quasi-likelihood F-test (glmQLFTest) was applied. To reduce noise from low expressed genes, only genes with CPM > 1 in at least three samples were included for analysis. Benjamini-Hochberg false discovery rates (FDR) were generated with a significance threshold <0.05. For each experiment, we tested seven pairwise comparisons (extracted from the factorial model): 1) acid/NO· stress vs TSB in WT, 2) acid/NO· stress vs TSB in ΔyqeK, 3) ΔyqeK vs. WT in TSB, 4) ΔyqeK vs. WT in acid/NO· stress, 5) interaction NO·/acid (whether stress response differs between genotypes), 6) condition effect for NO·/acid (main effect of stress across genotypes), and 7) genotype effect (main effect of yqeK deletion across conditions). For the first four pairwise comparisons the log2 fold change was calculated as logFC_total = logFC_genotype + logFC_interaction (e.g. WT vs ΔyqeK in acid or NO·) or logFC_total = logFC_condition + logFC_interaction (e.g. TSB vs NO· in yqeK). The acid stress batch TSB libraries were used for the primary Δ*yqeK* vs. WT comparison as they yielded greater statistical power; results from the NO· stress batch libraries were qualitatively consistent. Genes were assigned to functional categories based on keyword matching from gene annotations retrieved from the NCBI Datasets gene table and the Gene Specific Information table from AureoWiki (54) for USA300_FPR3757, followed by manual review; category assignments are listed in Table S3.

### Quantitative real-time PCR

For validation of RNA-Seq results, qRT-PCR was performed on RNA collected from three additional biological replicates of each strain during growth in TSB and acidified (pH 5.5) TSB. Primers are listed in Table S1. The *Power* SYBR™ Green RNA-to-C_T_™ *1-Step* Kit (Applied Biosystems) was used and reactions were performed on an Applied Biosystems StepOne Plus Real-Time PCR system. The housekeeping gene *rpoD* was used for normalization (55), and fold changes were calculated using the ΔΔCt method (56) with the average of three technical replicates for each reaction. Technical replicates within 1.5 Ct from the group median were retained for analysis, and qRT-PCR was repeated if there were fewer than two passing technical replicates for a sample.

### Hemolysis Assay

A quantitative hemolysis assay was performed using sheep red blood cells (RBCs). To prepare the RBCs, 1mL of sheep blood (Carolina Biological) was centrifuged at 1000xg for 5 minutes at 4℃ and washed 3x in cold PBS (with gentle inversion to avoid mechanical lysis). RBC pellets were weighed and resuspended to create a 2% RBC solution in PBS. The suspension was mixed by gentle inversion and kept on ice for same day use. Bacterial supernatants were prepared from 18-hour overnight cultures grown to similar optical densities (∼7) in TSB at 37°C; 1mL of overnight culture was pelleted and the supernatant was filter-sterilized with 0.22µm syringe filters to remove residual cells. Next, culture supernatants were mixed 50:50 with the 2% RBC suspension (500µL total volume), gently mixed by inversion, and incubated at 37°C for 1-hour without shaking (to avoid mechanical lysis). Water and PBS were used for positive and negative controls, respectively. After incubation, samples were centrifuged at 1000xg at 4°C for 5 minutes to pellet remaining RBCs. Supernatants (200µL) were transferred to 96-well plates, and absorbance at 540nm was quantified (average of three reads) to assess the amount of hemoglobin released. The percent hemolysis was determined by the formula % hemolysis=(sample-negative control)/(positive control-negative control)*100. A qualitative hemolysis assay was also performed by streaking a line of each strain on 5% sheep blood TSA plates and incubating at 37°C or 43°C for 48 hours.

## Statistical Analysis and Data Visualization

Statistical analyses and data visualization were performed in R (version 4.5.2; R Core Team, 2025) (57). For all assays, biological replicates represent independent experiments performed on separate days; technical replicates within biological replicates were averaged prior to statistical analysis. Differential expression analysis was performed with edgeR as described above. Growth curve nadir ratios, maximum growth rates, and hemolytic activity were compared across strains using one-way ANOVA followed by Tukey’s HSD post hoc test for pairwise comparisons. Luminescence data were analyzed by two-way ANOVA to account for both condition and strain, followed by Tukey’s HSD post hoc test for pairwise comparisons. Concordance between RNA-Seq and qRT-PCR fold changes was analyzed using Pearson correlation. Figures were created using ggplot2 (58). Cowplot (59) and Inkscape (1.3.2) were used to assemble figure panels.

## Acknowledgments

This work was supported by an AIREA grant from the American Heart Association (23AIREA1053493), funding from the UNC Asheville Undergraduate Research Program, the UNC Asheville Steve and Frosene Zeis Professorship, and the Forrest Fund for Undergraduate Research. J.V. was supported by a Chemistry Scholars Program Early Undergraduate Research Fellowship from National Science Foundation S-STEM grant 1833604 awarded to UNC Asheville. The funders had no role in study design, data collection, interpretation, or the decision to submit for publication. We are grateful to Dr. Anthony Richardson (Department of Microbiology and Molecular Genetics, University of Pittsburgh) for providing the WT *S. aureus* strains and allelic exchange vector pBT2*ts* for this study.

## Data Availability

Raw sequencing reads and processed count data have been deposited in the NCBI Gene Expression Omnibus (GEO) under accession numbers GSE319508 and GSE319509.

