## Supplemental Material for "Loss of the Ap_4_A hydrolase YqeK impairs stress adaptation and virulence gene expression in *Staphylococcus aureus*"

**Table S1.** Strains, plasmids, and primers.

| Strain | Description | Reference/Source |
|---|---|---|
| *S. aureus* RN4220 | Methicillin-Sensitive Restriction-Deficient *S. aureus* | A. Richardson |
| *S. aureus* LAC | Methicillin-Resistant Clinical *S. aureus* Isolate | A. Richardson |
| MG148 | S. aureus LAC Δ*yqeK* (SAUSA300_1552) | This study |
| MG011 | S. aureus LAC + pRMC2 | This study |
| MG162 | S. aureus LAC + pRMC2_yqeK | This study |
| MG166 | S. aureus LAC Δ*yqeK* + pRMC2 | This study |
| MG164 | S. aureus LAC Δ*yqeK* + pRMC2_yqeK | This study |
| **Plasmid** | **Description** | **Reference/Source** |
| pBT2*ts* | *E. coli*/*S. aureus* shuttle vector | (1) |
| pRMC2 | *S. aureus* expression vector | Addgene plasmid #68940 (2) |
| pBT2_yqeK | 5' and 3' homology regions of *yqeK* cloned into the BamHI site of pBT2ts to yield Δ*yqeK* | This study |
| pRMC2_yqeK | Complete *yqeK* allele with added Shine-Dalgarno cloned into the KpnI/EcoRI sites of pRMC2 | This study |
| **Primer** | **Sequence** | **Use/Source** |
| p314 | CACTA**GGATCC**CCTATTTTAGAGGAAGCAGAGC | *yqeK* deletion (this study) |
| p315 | GCTAATAATTCTTGTGAATTCATTTACATATAATCCTTCCCCCTTAAT | *yqeK* deletion (this study) |
| p316 | ATTAAGGGGGAAGGATTATATGTAAATGAATTCACAAGAATTATTAGC | *yqeK* deletion (this study) |
| p317 | CACTA**GGATCC**CTGCATCTTCATTATGTTCATC | *yqeK* deletion (this study) |
| p360 | CACTA**GGTACC**TACCCGGAGGAGATATTATGAACATTGAAAAGCAAAACGG | *yqeK* complementation (this study) |

| | | |
|---|---|---|
| p318 | CACTA**GAATTC**TTAATCATCCTTTATTCTTTCGTCAC | *yqeK* complementation (this study) |
| p369 | GATACGTAGGTCGTGGTATG | *rpoD* qRT-PCR (3) |
| p370 | CACGAGTGATTGCTTGTC | *rpoD* qRT-PCR (3) |
| p371 | CGATTAGGGATGCAGGTCTT | *agrB* qRT-PCR (this study) |
| p372 | CACCATGTGCATGTCTTCTT | *agrB* qRT-PCR (this study) |
| p373 | CAAGAAGCGATTGACCACAG | *tig* qRT-PCR (this study) |
| p374 | GCTTGTCCACCTTCGAATTC | *tig* qRT-PCR (this study) |
| p375 | GCGAATCTTGCAAGACTTCA | *gatC* qRT-PCR (this study) |
| p376 | AGGTTCAACGCCTTCTGTAT | *gatC* qRT-PCR (this study) |
| p377 | TGGCGTTGTTGGATATGGTA | *spn* qRT-PCR (this study) |
| p378 | TGTCTTTTGTAGCTTGGTCAG | *spn* qRT-PCR (this study) |
| p379 | GCGTCATTAGCGGGTATTTC | *cap8C* qRT-PCR (this study) |
| p380 | TGTGTTATGCGCATCTGAAC | *cap8C* qRT-PCR (this study) |
| p381 | TTTCACCAGACTTCGCTACA | *hla* qRT-PCR (this study) |
| p382 | TCCAGTGCAATTGGTAGTCA | *hla* qRT-PCR (this study) |
| p383 | TGGCACAAGATATCATTTCAACA | *hld* qRT-PCR (this study) |
| p384 | TGAATTTGTTCACTGTGTCGA | *hld* qRT-PCR (this study) |
| p385 | TAGTGGTTTAGCTCAAGGGC | *hrtB* qRT-PCR (this study) |
| p386 | CTCAATTTGCGGCTCTTTCA | *hrtB* qRT-PCR (this study) |

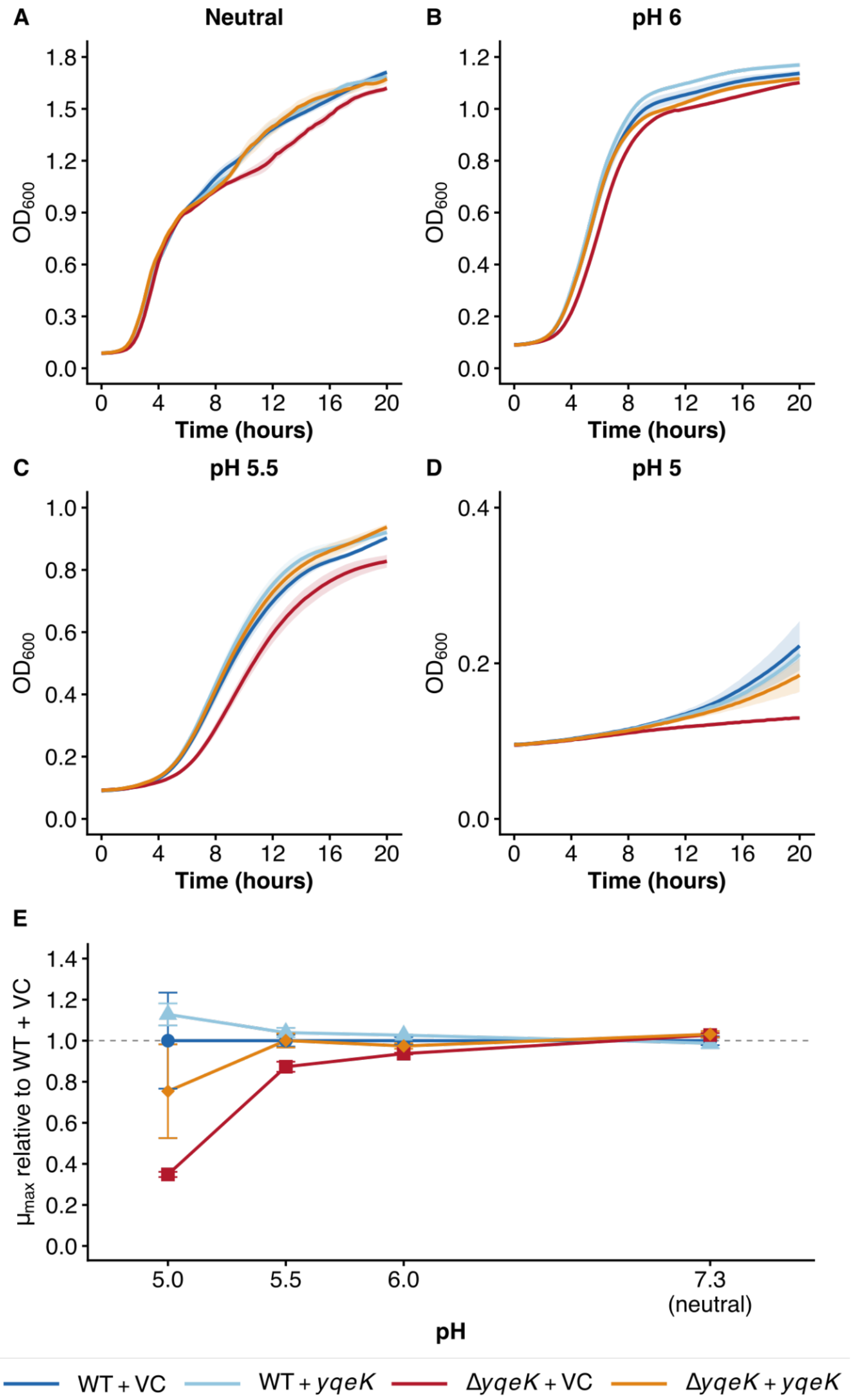


**Figure S1. Δ*yqeK* exhibits a larger growth defect relative to WT as pH decreases.** Growth curves of the *S. aureus* LAC strains WT + pRMC2 (vector control, VC), Δ*yqeK* + VC, WT + pRMC2-yqeK, and Δ*yqeK* + pRMC2-yqeK (complemented mutant) were performed in 96-well plates either in A) regular TSB (pH ~7.3) or TSB acidified with acetic acid to B) pH 6, C) pH 5.5, or D) pH 5. Growth curves are plotted as means; shaded regions represent SEM. E) Maximum specific growth rates ($\mu_{max}$) were calculated and plotted as a percentage of the $\mu_{max}$ of WT + VC under each condition.

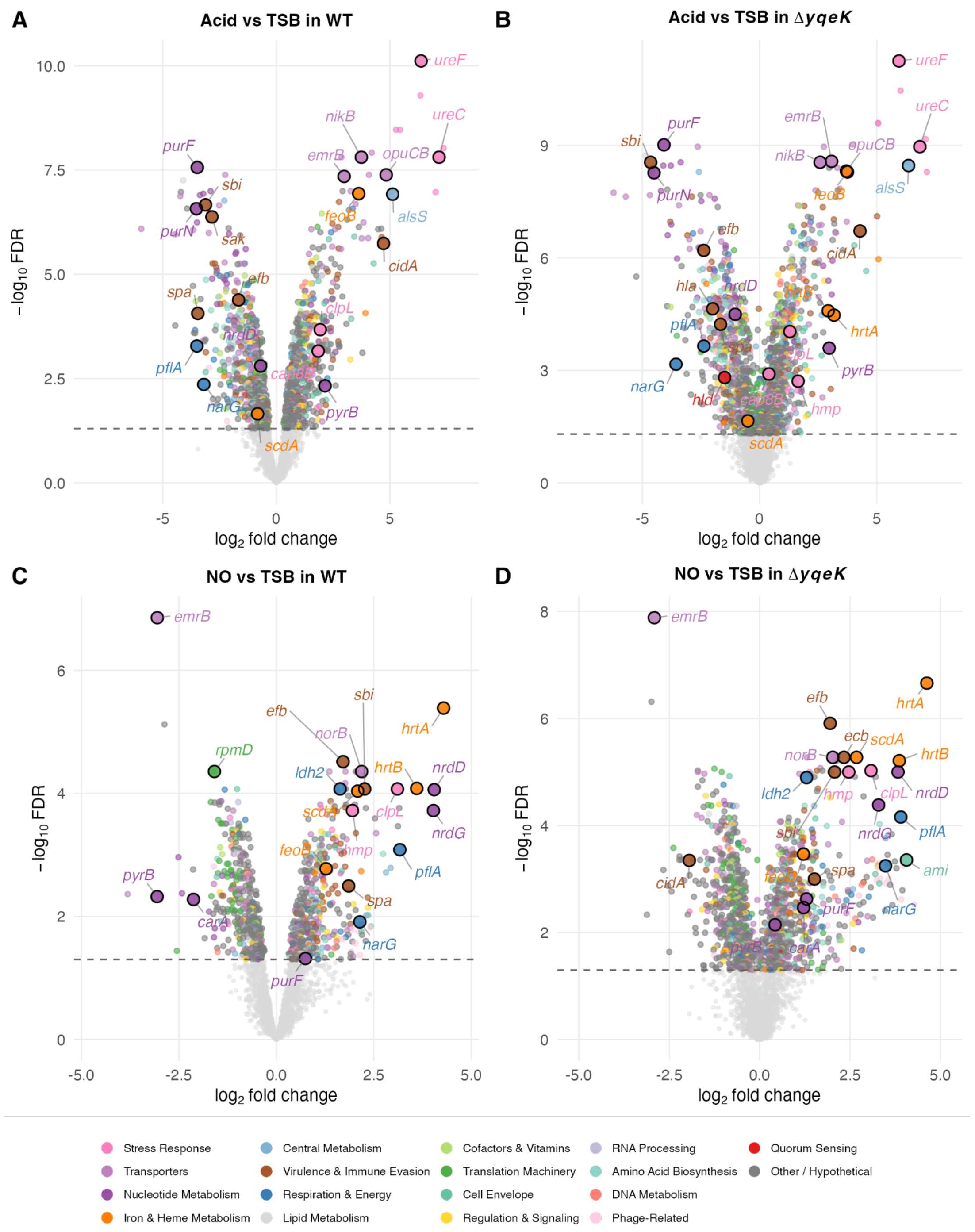


**Figure S2. Transcriptome changes within each strain across conditions.** Volcano plots showing differential gene expression between growth in acid stress vs. TSB for A) WT and B) Δ*yqeK*, as well as NO· stress vs. TSB for C) WT and D) Δ*yqeK*. Genes were labeled based on a combined significance score ($|\log_2 FC| \times -\log_{10} FDR$) to highlight the most statistically robust differentially expressed genes. Labeled genes were selected from those with annotated functions; hypothetical proteins are plotted but not labeled for clarity.
